# Genomic insights of high-risk clones of ESBL-producing *Escherichia coli* isolated from community infections and commercial meat in Southern Brazil

**DOI:** 10.1101/2020.12.31.424884

**Authors:** João Gabriel Material Soncini, Louise Cerdeira, Vanessa Lumi Koga, Ariane Tiemy Tizura, Gerson Nakazato, Renata Katsuko Takayama Kobayashi, Caio Augusto Martins Aires, Nilton Lincopan, Eliana Carolina Vespero

## Abstract

During a microbiological and genomic surveillance study to investigate the molecular epidemiology of extended-spectrum beta-lactamase (ESBL)-producing *Escherichia coli* from community-acquired urinary tract infections (UTI) and commercial meat samples, in a Brazilian city with a high occurrence of infections by ESBL-producing bacteria, we have identified the presence of CTX-M (−55, −27, −24, −15, −14 and −2)-producing *E. coli* belonging to the international clones ST354, ST131, ST117, and ST38. The ST131 was more prevalent in human samples, and worryingly the high-risk ST131-C1-M27 was identified in human infections for the first time. We also detected CTX-M-55-producing *E. coli* ST117 isolates from meat samples (i.e., chicken and pork) and human infections. Moreover, we have identified the important clone CTX-M-24-positive *E. coli* ST354 from human samples in Brazil for the first time. In brief, our results suggest a potential of commercialized meat as a reservoir of high-priority *E. coli* lineages in the community. In contrast, the identification of *E. coli* ST131-C1-M27 indicates that novel pandemic clones have emerged in Brazil, constituting a public health issue.

## INTRODUCTION

*Escherichia coli* is a commensal of the human intestinal tract and most warm-blooded mammals and figures as an important pathogen for humans and animals^1,8^. In humans, urinary tract infection (UTI) is the second most common bacterial infection managed in primary care, and uropathogenic *E. coli* (UPEC) is responsible for 75% to 95% of the cases^1^. The increasing antimicrobial resistance (AMR) detected in clinical UPEC isolates has been of concern^1^ and infections caused by antimicrobial-resistant bacteria as extended-spectrum β-lactamase (ESBL)-producing *E. coli* represent significant healthcare issues^8^ since it compromises the effective treatment, being responsible for a large number of morbidity and mortality^4^.

Since *E. coli* can act as a large reservoir of resistance genes that directly impact treatment in human and veterinary medicine, the debate over the transmission of multiresistant *E. coli* strains between animals and humans through numerous pathways has become increasingly important. However, the interaction between food-producing animals, humans, and the environment regarding the transmission of these resistant pathogens is not yet fully understood^8,9^.

The isolation of ESBL-producing *E. coli* from food-production animals is increased worldwide, mostly from chicken meat^8,9^. The excessive use of antimicrobials in livestock is one of the practices that help in the emergence of pathogens resistant to humans. The consumption of meat, direct contact with colonized animals, or manure spread in the environment are sources for the transmission of livestock AMR to humans^5,6^. Besides that, AMR gene transfer may occur between different bacterial species in the gut of animals and humans^7^.

The CTX-M type is one of the largest groups of ESBL, and recent studies that addressed the epidemiology of these enzymes in Brazil, show that CTX-M-2, CTX-M-8, CTX-M-9, and CTX-M-15 are the predominant variants in the country^10–12^. Many types of CTX-M-producing *E. coli* have been recognized as belonging to specific clones commonly isolated from UTI cases originating in a particular locale, country, or even globally. Some studies show that isolates from foods CTX-M genotypes sometimes correspond with the locally dominant human types^13,14^.

Considering the emerging AMR in Brazil, both in human medicine as in livestock, and the need for understanding this panorama, we conducted next-generation sequencing (NGS)-based analysis adopting a One Health approach to assess national transmission of CTX-M-producing *E. coli* isolated from meat products and human patients.

## RESULTS

The results for each of 91 *E. coli* isolates included in this study can be seen in **Figure 1.** It is notorious high rates of resistance to ampicillin (100%), ceftriaxone (87.91%), nalidixic acid (87.91%), cefepime (83.52%), trimethoprim-sulfamethoxazole (82.42 %), nitrofurantoin (76.92%), norfloxacin (75.82%) and ciprofloxacin (72.53%). Less than half showed resistance to gentamicin (36.26%) and amoxicillin/clavulanate (21.98%). Only three (3.30%) isolates were resistant to piperacillin-tazobactam and two (2.20%) to amikacin. Some isolates also showed intermediate resistance levels: 28.57% to amoxicillin/clavulanate; 4.40% to piperacillin-tazobactam and gentamicin; 1.10% to ciprofloxacin and norfloxacin.

**Figure 01.**
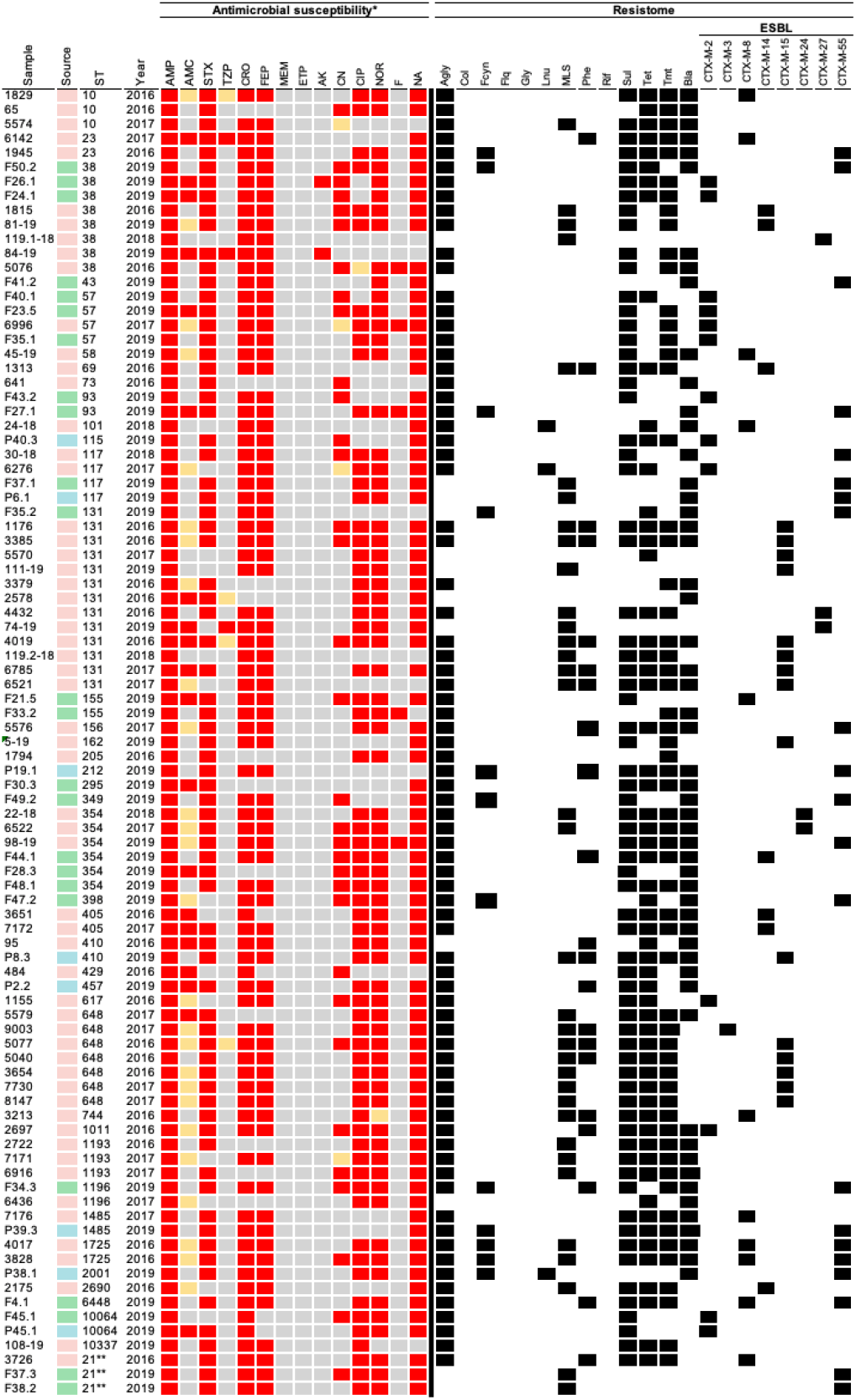
Heatmap show AMR profile, ST and Resistome.

Genomic analysis revealed 57 genes associated with resistance to aminoglycosides (n = 15), β-lactams (n = 12), trimethoprim (n = 8), phenicols (n = 5), tetracyclines (n = 4), macrolides (n = 4), sulfonamides (n = 3), quinolones (n = 3), lincosamides (n = 2) and fosfomycin (n = 1). Regarding aminoglycosides, the most prevalent genes were *strA* and *strB,* both with 39.56%, followed by the *aadA1* gene (36.26%). The *dfrA17* and *dfrA1* genes, associated with resistance to trimethoprim, were detected in 31 (34.07%) and 13 (14.29%) isolates, respectively. Genes related to phenicols resistance had similar prevalence, being *catB3* (8.79%), *floR* (5.49%), *catA1* (4.40%) and *cmlA1* (4.40%). Concerning to tetracyclines resistance, we detected the *tet(A)* (38.46%) and *tet(B)* (27.47%) genes. About macrolides the *mph(A)* gene (29.67%) was found and the detected genes related to sulfonamides were *sul1* (56.04%) and *sul2* (53.85%). Few isolates had lincosamide resistance genes, two of them had *Inu(F)* (2.20%) and one *Inu(A)* (1.10%). The *fosA* gene found in three isolates was the only one associated with fosfomycin resistance.

The genes associated with resistance to β-lactams were *bla*_TEM-1B_ (48.35%), *bla*_OXA-1_ (7.69%), *bla*_CMY-2_ (6.59%), and *bla*_TEM-1A_ (2.20%), in addition to eight variants of the *bla*_CTX-M_ gene that encode CTX-M-type ESBL enzymes. Among the ESBL coding genes, *bla*_CTX-M-55_ was the most detected (21.98%), mainly from chicken meats (n = 10), followed by humans (n = 6) and porks (n = 4). The *bla*_CTX-M-15_ was found predominantly in human isolates (n = 14) and only in one pork isolate. On the other hand, *bla*_CTX-M-2_ was also detected in 15 isolates (16.48%), being them chicken meat (n = 7), human (n = 6) and pork (n = 2). The CTX-M-8 and CTX-M-14 coding genes were present in eight and five human isolates, and two and one chicken meat isolates, respectively. The CTX-M-24 (n = 4), CTX-M27 (n = 3) and CTX-M-3 (n = 1) coding genes were present only in human isolates.

In this work, 52 plasmid incompatibility groups belong to the p0111, IncF, IncI1, and IncN families. In human isolates, the most frequent pMLST were IncI1[ST-113] (n=9), IncF[F-: A-: B-] (n=7), IncF[F1: A2: B20] (n=5), IncF[F48: A1: B49] (n=5) and p0111 (n=5). In chicken meat isolates, IncF[F18: A-: B1] (n=8), p0111 (n=7) and IncN [Unknown ST] (n=5) were the most frequent pMLST. In isolates of pork, the most frequent incompatibility groups were IncN[Unknown ST] (n=4) and IncF[F33: A-: B1] (n=3).

In total, 40 sequence types (STs) were found, the most observed were the ST131 (n = 12), ST38 (n = 8), ST648 (n = 7), and ST354 (n = 6). Some STs were detected in more than one source, demonstrating a genetic relationship between these isolates, mainly between humans and chicken meat. The ST38, ST131, ST354, and ST1196 were found in both urine and chicken meat strains in the respective quantities of 5 and 3, 12 and 1, 4 and 3 and 1 and 1. The ST410 was the only observed in urine (n = 1) and pork (n = 1) strains. The ST117 was present in the three sources studied, with two strains from urine, one from chicken meat and pork. The clonal relationship between the isolates in this study and the dissemination distribution in Brazil can be seen in **Figure 2A-C**. Additionally, it is possible observed that UK sample (ST131) clustered with other samples isolates in Brazil (ST131), all results could be view in Microreact link (https://microreact.org/project/2mKg54AHdWj5xdJ5VFejY8).

**Figure 02.**
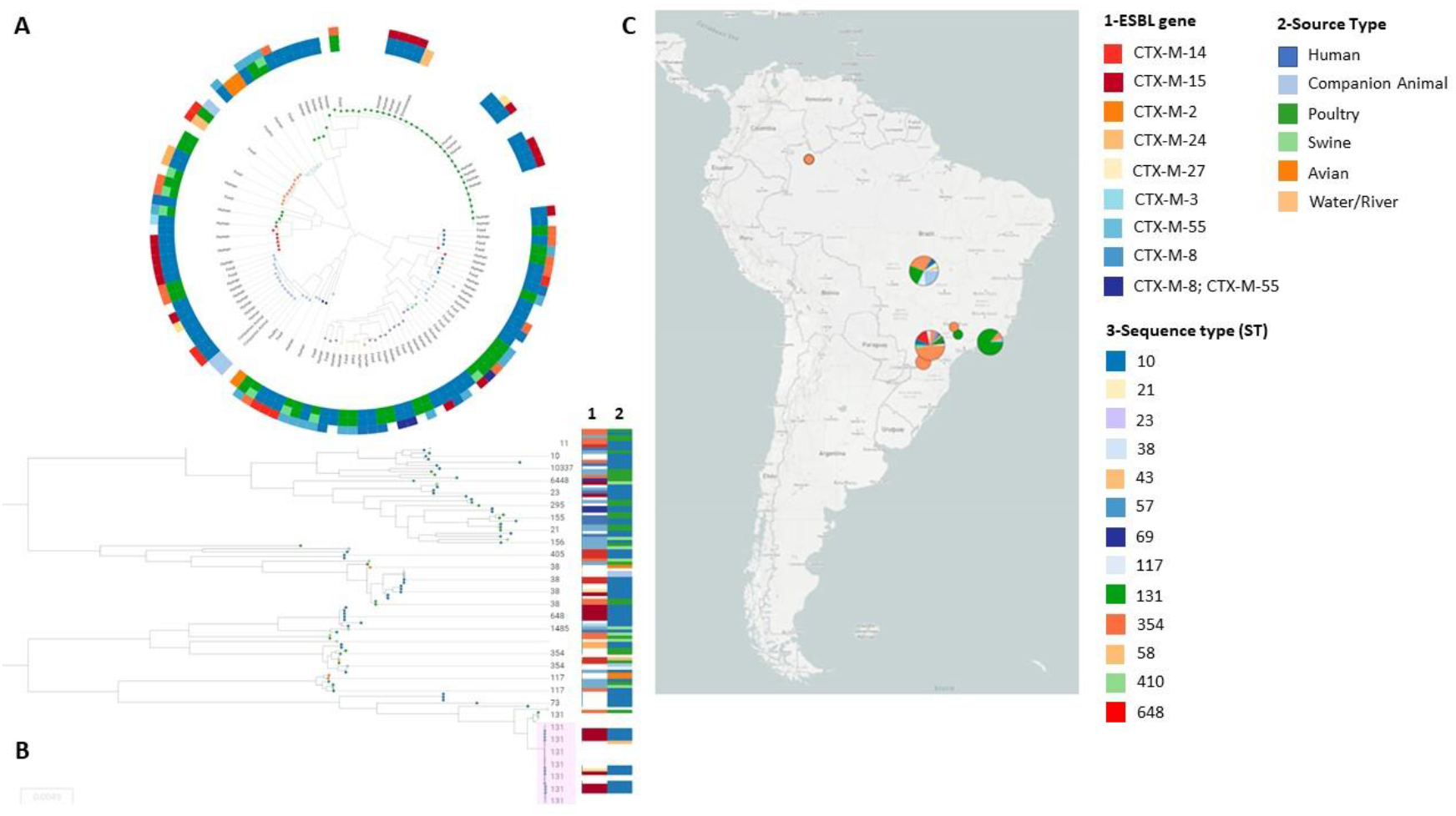
(A) *E. coli* Phylogenomic SNP tree with circular heatmap shows source type (first inner circle), sequence type (second inner circle) and ESBL-gene (third inner circle. (B) *E. coli* Phylogenomic SNP tree with columns shows ESBL-gene (column 1) and source type (column 2); light pink highlights the clade where UK-ST131 samples clustered with Brazilian ST131 samples. (C) dissemination distribution map in Brazil.

## DISCUSSION

This study presents the first reports of *E. coli* ST131-C1-M27 in human infection and CTX-M-24-positive *E. coli* ST354 from ITU, in Brazil. In Latin America CTX-M-producing *E. coli* are endemic. Our data show a wide distribution of these isolates belonging to the international clones in livestock and the community. The extensive presence of CTX-M enzyme-producing strains in several sources raises the hypothesis that the spread occurs with greater frequency and efficiency, especially among enterobacteria^10^.

*E. coli* ST131 globally known and is related to the spread of resistance genes, including specific CTX-M coding genes^15^. Recent studies have shown that ST131 is rare among animal isolates, becoming almost exclusively a human pathogen, as demonstrated by our results, where ST131 is predominantly found in strains of human urine^16^. The subclade C2 is associated with *bla*_CTX-M-15_ that can be carried by different groups of plasmids^17^. Here we also observe that all *bla*_CTX-M-15_ are involved with the incompatibility group IncF. In a study by Peirano et al. (2020), it was shown that clade C was related to the highest rates of UTI, with subclade C2 being the most common and associated with incompatibility group IncFII^18^. Besides, CTX-M-15-producing *E. coli* ST131 has already been shown to be involved in outbreaks in health institutions and is the most prevalent ESBL-producing *E. coli* worldwide^19^.

The CTX-M-27-producing ST131-C1 has been considered a new epidemic clone, and there have been no reports of human infections so far, in Brazil. Clade C1-M27 is associated with CTX-M-27 and was first observed as colonizing children in France in 2012. Recent studies suggest that the subclade C1-M27 was recently selected since SNPs have a smaller difference between isolates of this same subclade than SNPs of isolates of subclade C2 and A. In addition, the plasmid predominantly involved with the dissemination of *bla*_CTX-M-27_ is IncF[F1:A2:B20], as found in our study. Resistance to fluoroquinolones, macrolides, tetracyclines, aminoglycosides, and sulfonamides appears to be part of the profile of C1-M27 isolates^20,21^.

The CTX-M-14 and CTX-M-24 enzymes belong to the CTX-M-9 group. Although the first one is widely distributed worldwide, especially in China, South-East Asia, Japan, South Korea, and Spain, microorganisms producing CTX-M-24 remain relatively rare, reported with greater incidence in countries such as Peru and Bolivia^18,22,23^. This study found an important association between CTX-M-24 and *E. coli* ST354 detected in two human isolates, never before reported in UTI in Brazil. In a study by Dagher et al. (2018), ST354 isolates were positive to *bla*_CTX-M-24_ and resistant to ciprofloxacin, associated with extra-intestinal infections, animals and humans, reinforcing the zooanthroponotic hypothesis of these clones^24^.

ESBL type CTX-M-2 and CTX-M-55 are frequently found, and their coding genes are spread in several ways. Some studies suggest that the plasmid IncF[F33: A-: B-] is involved in disseminating these genes, which may explain the coexistence of these two genes in two strains belonging to ST1725 isolated from urine samples^25^. Although another strain of *E. coli* ST6448 isolated from chicken meat also showed the coexistence of *bla*_CTX-M-2_ and *bla*_CTX-M-5_, the plasmids that carried them belonged to IncF [F24: A-: B73] and IncI1 [ST −131], respectively. In the last ten years, IncI1-type plasmids have had a high spread, mainly in animal reservoirs. There are reports of *bla*_CTX-M-2_, *bla*_CTX-M-8_ and *bla*_CTX-M-55_ genes frequently found on IncI plasmids from *E. coli* isolated from chickens and pigs several countries, such as China, France, the United States of America, and the United Kingdom^26–28^.

The international clone ST117, found in the three different sources of this study, is often found in chicken meats and pork, and it is also associated with human infections. Studies have already reported the multiple resistance profile of ST117 and associated it with CTX-M-55 expression, consistent with our results^29,30^. Likewise, ST38 is also widely found in chickens and humans, worldwide, and is related to several ESBL genes, such as *bla*_CTX-M-14_, *bla*_CTX-M-27_, and *bla*_CTX-M-55_. One of the hypotheses for the successful dissemination of these genes among the *E. coli* clones is that the families of plasmids IncI1 and IncF are important vectors for disseminating *bla*_CTX-M_. In China, South Korea, and Japan, studies suggest an epidemic of *bla*_CTX-M_ genes carried by plasmids IncI1, IncF[F33: A-: B-], IncF[F46: A-: B20] and IncF[F18: A-: B1], found in cattle, pigs, chickens, pets and humans^31–33^. The second hypothesis suggests that *E. coli* ST131 isolated in UK in 2001, could be the origin clone of *E. coli* ST131 disseminated in Brazil, and may be after arrived in Brazil this clone acquired a plasmid carrying *bla*_CTX-M-55_ gene.

In conclusion, *E. coli* carrying *bla*_CTX-M_ genes from different sources seem to be related to the spread of internationally known clones (ST354, ST131, ST117, ST38). Some clones associated with some CTX-M variants are more prevalent in some sources than others do not exclude the possibility that new clones are entering and establishing themselves in different niches, as shown in this study. Thus, novel studies should continue to be carried out with more samples and sources to understand further the dynamics of dissemination, shift, and establishment of ESBL-producing *E. coli* clones at the interface between animal sources and human health.

## MATERIAL AND METHODS

### Study population

During June 2016 to May 2019, 195.080 urine cultures were performed in a by public health services, in a city in south of Brazil. A total of 34.293 (17,6%) were positive for gram-positive or gram-negative microorganisms; of these 22.698 (66,2%) were E. coli strains and a total of 2.033 (6,2%) ESBL producing bacteria, being 1.389 (51,2%) ESBL production *E. coli.* Concomitantly, a surveillance study from January to May 2019 was carried out, to research ESBL-producing *E. coli,* in chicken and pork meat, bought at markets and butcher shop near public health services. Fluoroquinolone-resistant and ESBL-producing *E. coli* were investigated in chicken meat (n = 50), and pork (n = 50) samples. A total of 102 *E. coli* was isolated from chicken meat marketed, with 52 ESBL positive. And 67 resistant *E. coli* were isolated in pigs, 31 ESBL positive. This study included for sequencing 91 *E. coli* strains of 102 total isolates: 59 isolated from urine culture(n=59), chicken meat (n=24) and pork (n=8). These strains were selected by the similarity profile established by ERIC-PCR analysis of 1.389 ESBL-producing isolates. The study was approved by the Ethics and Research Committee of the State University of Londrina CAAE 56869816.0.0000.5231.

### Microbiological methods

Urine collected from women patients was inoculated on CHROMagar (Becton Dickinson, Heidelberg, Germany) and MacConkey (Merck, Darmstadt, Germany) plates using a calibrated inoculating loop with a capacity of 10 μl and incubated at 37°C for 24h.

The samples of chicken meat and pork were dipped in Brain Heart Infusion broth (Oxoid) with cefotaxime (4μg / mL), ciprofloxacin (4μg / mL), and both (Sigma-Aldrich, Munich, Germany) to selected resistant *E. coli* strains. After incubation, the solution was inoculated in the same way used for urine samples. All the isolates were stored in Tryptic Soy Broth (TSB) with 15% glycerol (−20°C).

The identification and bacterial susceptibility were performed by the automated VITEK^®^ 2 system, using the VITEK^®^ 2 AST 239 card and the VITEK^®^ 2 GN ID card (BioMérieux, USA). The bacterial susceptibility was tested for 14 antibiotics: ampicillin, amoxicillin/clavulanate, ceftriaxone, cefepime, ertapenem, meropenem, nalidixic acid, ciprofloxacin, norfloxacin, gentamicin, amikacin nitrofurantoin, trimethoprim-sulfamethoxazole, and piperacillin-tazobactam. The CLSI 2020 (Clinical and Laboratory Standards Institute) criteria were used for interpretation. *E. coli* ATCC^®^25922 strain was used as quality control.

### ERIC-PCR

1.389 ESBL-producing isolates were subjected to Enterobacterial Repetitive Intergenic Consensus (ERIC-PCR), by Versalovic et al. (1991)^34^. Analysis of genomic fingerprinting was performed using GelJ v.2.0 software by the Dice similarity method (HERAS et al., 2015)^35^. Strains were considered genetically related if the similarity index was ≥85 %.

### DNA isolation and whole-genome sequencing

For DNA extraction, strains were grown on Mueller-Hinton Agar overnight at 37 °C. Subsequently, a single colony was inoculated in 2 mL of Luria-Bertani broth for 12 hours at 37 °C. The suspension was used to continue extraction and purification by the DNA extraction kit (Invitrogen, Carlsbad, CA). The extracted DNA was quantified by Qubit dsDNA (double-stranded DNA) BR assay kit (Invitrogen, Carlsbad, CA). After quantification, the DNA was used to construct a paired-end library (150 bp), sequenced using the NextSeq platform (Illumina). The instructions of each manufacturer were followed in all steps.

### Bioinformatic analysis

Genome quality filter and assemblies were performed by the CLC Genomics Workbench version 7.0 (Aarhus, Denmark). Multilocus sequence type (MLST), resistome, and virulome were identified using MLST v2.0 (Larsen et al., 2012), ResFinder v3.1(Bortolaia et al., 2020), VirulenceFinder v2.0, (Joensen et al., 2014), PlasmidFinder v2.1 (Carattoli et al., 2014), FimTyper v1.0 (Roer et al., 2017) and SerotypeFinder v.2.0 (Joensen et al., 2015), respectively. The BacMet database (Pal et al., 2013) was used to identify biocides and heavy metal (HM)^31,36–42^. The EnteroBase (https://enterobase.warwick.ac.uk/) was used to create a single nucleotide polymorphisms (SNPs) project to strains that showed the same STs genomes were aligned against genomes of other Brazilian studies.

## ACKNOWLEDGEMENTS

The authors would like to thank the financial support of the Bill and Melinda Gates Foundation’s Grand Challenges Explorations Brazil – New Approaches to characterize the global burden of antimicrobial resistance (OPP1193112), the Research Support Facilities Center of the State University of São Paulo (CEFAP-USP), Institute of Biomedical Sciences of State University of São Paulo (ICB-USP) and Master’s program Clinical and Laboratory Pathophysiology of the State University of Londrina.

## AUTHOR INFORMATION

### Affiliations

*Laboratory of Clinical Microbiology, Department of Pathology, Clinical and Toxicological Analysis, Health Sciences Center, State University of Londrina, Londrina, Brazil*

João Gabriel Material Soncini

*Department of Infectious Diseases, Central Clinical School, Monash University, Melbourne, Australia*

*Department of Vector Biology, Liverpool School of Tropical Medicine, Liverpool, United Kingdom.*

Louise Cerdeira

*Department of Microbiology, Biological Science Center, State University of Londrina, Londrina, Brazil*

Vanessa Lumi Koga

*Laboratory of Clinical Microbiology, Department of Pathology, Clinical and Toxicological Analysis, Health Sciences Center, State University of Londrina, Londrina, Brazil*

Ariane Tiemy Tizura

*Department of Microbiology, Biological Science Center, State University of Londrina, Londrina, Brazil*

Gerson Nakazato

*Department of Microbiology, Biological Science Center, State University of Londrina, Londrina, Brazil*

Renata Katsuko Takayama Kobayashi

*Department of Health Science, Health and Biological Science Center, Federal Rural University of Semi-Arid, Mossoró, Brazil.*

Caio Augusto Martins Aires

*Department of Microbiology, Institute of Biomedical Sciences, University of São Paulo, Brazil*

Nilton Lincopan

*Laboratory of Clinical Microbiology, Department of Pathology, Clinical and Toxicological Analysis, Health Sciences Center, State University of Londrina, Londrina, Brazil*

Eliana Carolina Vespero

### Data availability

Draft whole-genome assembly was deposited in GenBank under the bioproject PRJNA578368. The data of the figures can be accessed in Figshare (https://doi.org/10.6084/m9.figshare.12808439.v1).

## ETHICS DECLARATIONS

### Ethical approval

The study was approved by the Ethics and Research Committee of the State University of Londrina CAAE 56869816.0.000.5231.

